# Federated Learning for Diagnosis of Age-related Macular Degeneration

**DOI:** 10.1101/2023.07.06.547937

**Authors:** Sina Gholami, Jennifer I. Lim, Theodore Leng, Sally Shin Yee Ong, Atalie Carina Thampson, Minhaj Nur Alam

## Abstract

This paper presents a federated learning (FL) approach to train deep learning models for classifying age-related macular degeneration (AMD) using optical coherence tomography image data. We employ the use of residual networks and vision transformer encoders for the normal vs AMD binary classification, integrating four unique domain adaptation techniques to address domain shift issues caused by heterogeneous data distribution in different institutions. Experimental results indicate that FL strategies can achieve competitive performance similar to centralized models even though each local model has access to a portion of the training data. Notably, Adaptive Personalization FL strategy stood out in our FL evaluations, consistently delivering high performance across all tests due to its additional local model. Furthermore, the study provides valuable insights into the efficacy of simpler architectures in image classification tasks, particularly in scenarios where data privacy and decentralization are critical using both encoders. It suggests future exploration into deeper models and other FL strategies for a more nuanced understanding of these models’ performance.

## 1 Introduction

Age-related macular degeneration (AMD) is a common eye condition and a leading cause of vision loss among people aged 50 and older [36]. AMD causes damage to the macula, the part of the eye that provides sharp, central vision, which is located near the center of the retina. As a result, everyday activities such as reading and driving may be difficult to perform. In order to prevent severe vision impairment and preserve vision, detection of AMD in its early stages are crucial to implementing appropriate treatments, such as medications or procedures. Artificial intelligence (AI) can play a pivotal role in the preliminary identification and classification of AMD [9, 4, 14, 27, 17, 39, 33]. Its proficiency in discerning the disparate stages of both wet and dry AMD results in substantial enhancement of the prognosis of treatment outcomes. Deep learning (DL) models significantly refine the precision and accuracy of AMD diagnosis, capable of detecting subtle ocular changes that might elude human scrutiny [5]. The remarkable capacity of AI for rapid analysis of imaging data facilitates more expeditious and efficient diagnosis, a critical factor in timely disease management [3].

The broad applications of AI include large-scale AMD screening within populations, a critical feature, particularly in areas where accessibility to ophthalmologists is restricted [26]. Beyond these clinical uses, AI’s potential to discern patterns and correlations in expansive datasets could yield innovative perspectives into the origins and evolution of AMD, potentially influencing future research trajectories [1]. AI models employed in AMD diagnosis predominantly utilize centralized learning. This traditional method accumulates data from diverse sources, collating them in a centralized server or location for the training of a machine learning (ML) model [37]. Adherence to data protection regulations such as the Health Insurance Portability and Accountability Act (HIPAA) is paramount in healthcare environments [32]. Thus, this approach encounters hurdles due to data privacy and security concerns in the medical sphere.

The introduction of federated learning (FL) allows model training without the dissemination of raw patient data, thereby circumventing privacy issues as data remains local. The possibilities proffered by FL involve enhancing diagnostic accuracy, prediction capability, and personalized treatment within ophthalmology, whilst harnessing large, diverse datasets from multiple institutions. However, for successful FL integration, it is necessary to address challenges linked with data heterogeneity, along with assuring the reliability and security of the learning process.

Inconsistencies in optical coherence tomography (OCT) image acquisition parameters and scanning protocols can induce variations in image quality [8]. Clinical and technical hurdles including differing standards and regulations among various institutional review boards, and limited training datasets for rare diseases can exacerbate the complexities of constructing and implementing DL techniques [31]. Such variations can impact the competence and generalizability of DL models [7].

The problem of domain shift also poses a significant challenge in the context of FL [21, 29]. Domain shift arises when there is a substantial difference in data distributions across various local devices or nodes, also termed as clients, within the FL system. The non-identically distributed nature of decentralized data, a key characteristic of FL, can potentially compromise model learning performance [19]. Rectifying this issue necessitates strategic and robust methodologies.

In the research of [19], domain adaptation (DA) techniques are outlined for optimizing learning algorithms irrespective of disparities in data distribution. Employing domain-invariant features or transfer learning methodologies, these techniques endeavor to lessen the impact of varied data distributions. Additionally, data augmentation can be leveraged to artificially enhance data representation, thereby diminishing the effects of domain shift [24]. An array of other methods can also be utilized to counter this challenge, encompassing client selection and sampling strategies, model aggregation procedures, proactive domain exploration [38], and FL personalization [35]. By effectively tackling domain shift, FL can bolster the model’s generalization capacity and augment performance across disparate domains.

This manuscript delineates the practicality of employing DA FL in the diagnosis of AMD. Using data from three distinct datasets, this study examines various FL strategies to address the domain shift issue for retinal OCT binary classification for AMD. The performance of these FL strategies is compared with a baseline centralized approach, emphasizing the potential benefits of employing multiple FL techniques to counteract the domain shift.

## 2 Methods

### 2.1 Data

We leveraged OCT data derived from three distinct research datasets for our study: [15], [30], [18], hereinafter referred to as DS1, DS2, and DS3. The utilization of these distinct datasets facilitated the simulation of three disparate institutions (FL nodes) intent on training a DL model for binary image classification (Normal Vs AMD). Hence, each node is allocated its own training, validation, and testing set.

DS1 encompasses a total of 84,484 OCT retinal images from 3,919 patients which are classified into four categories: Normal, Choroidal Neovascularization, Diabetic Macular Edema (DME), and Drusen. These images are compartmentalized into three separate folders: training, validation, and testing. However, it was observed that some images were duplicated across the validation and testing folders as well as the training folder. To eliminate redundancy, we amalgamated the validation and testing folders and compared each image with those in the training set using the mean square error (MSE) technique. An MSE score of zero signified the presence of identical images, leading to the identification of 8,520 duplicates within the dataset. These issues originated from 34 images that were marked as both normal and diseased retina. We discarded these images and exclusively used Drusen and Normal retinal images for this binary classification task. In the end, around 3% of the patient samples were chosen as the test set, which contained varying numbers of scans per patient.

DS2 comprises retinal images from 45 subjects, including 15 individuals each with Normal retinas, AMD, and DME. The initial 11 patients’ data was designated for training, the 12th for validation, and the remaining data were used for testing the model.

DS3 includes OCT imaging of 500 subjects captured under two fields of view: 3-mm and 6-mm. Each 3-mm file contains 304 scans from a single patient, while each 6-mm file accommodates 400 scans. Given the lack of a substantial role of peripheral retinal parts in classification, we focused only on fovea images (image numbers 100 to 180 for 3-mm and 160 to 240 for 6-mm).

The distribution of the data across the three datasets is visually represented in Figure 1 and tabulated in Table 1. The datasets for training, validation, and testing have been resized to a resolution of 128 × 128. To enhance the strength and ability to handle variations in different datasets, our DL networks have incorporated data augmentation techniques [25, 28]. These techniques involve random horizontal flipping, elastic transformations, and affine transformations.

**Figure 1:**
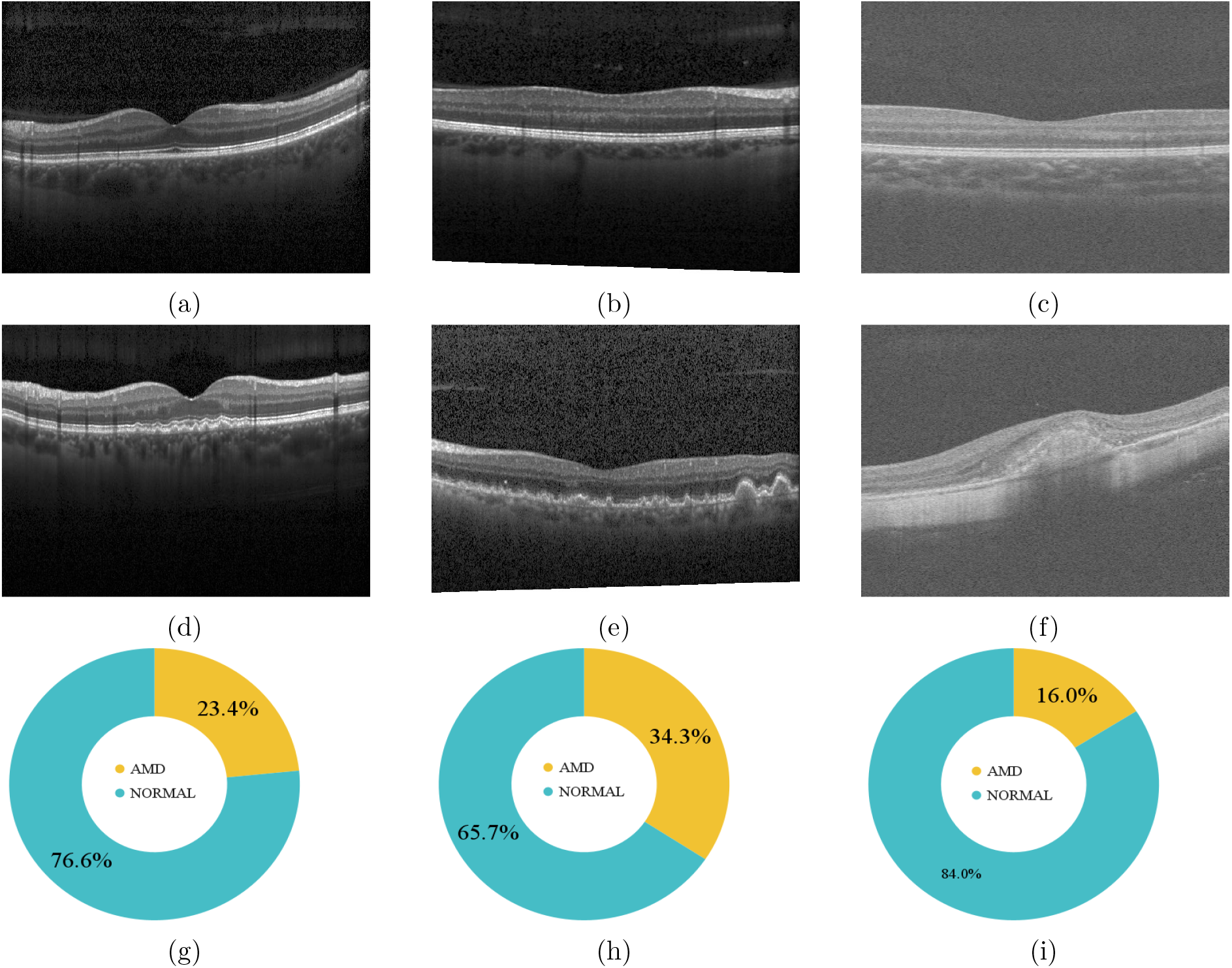
(**A**)-(**C**) Normal and (**D**)-(**F**) AMD sample OCTs from the three datasets. (**G**)-(**I**) illustrate how DS1, DS2, and DS3 are distributed.

**Table 1:**
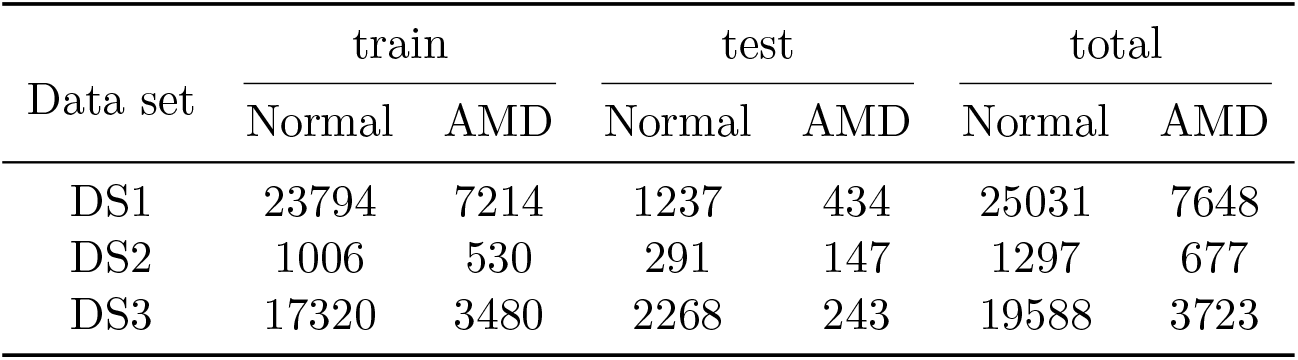
The data distribution of the three data sets for the training and test sets, including the number of normal and ADM retinas for each set.

To gain insights into the distribution of our datasets, which in turn would aid in evaluating the performance of our models across the test sets, we calculated the average histogram of all OCT images in each dataset. These histograms are visualized in Figure 2, providing a clear picture of the individual dataset distributions. Upon observation, it is evident that the distributions of DS1 and DS2 are quite similar. In contrast, DS3 displays a wholly distinct distribution, likely stemming from the unique imaging protocols utilized in its creation.

**Figure 2:**
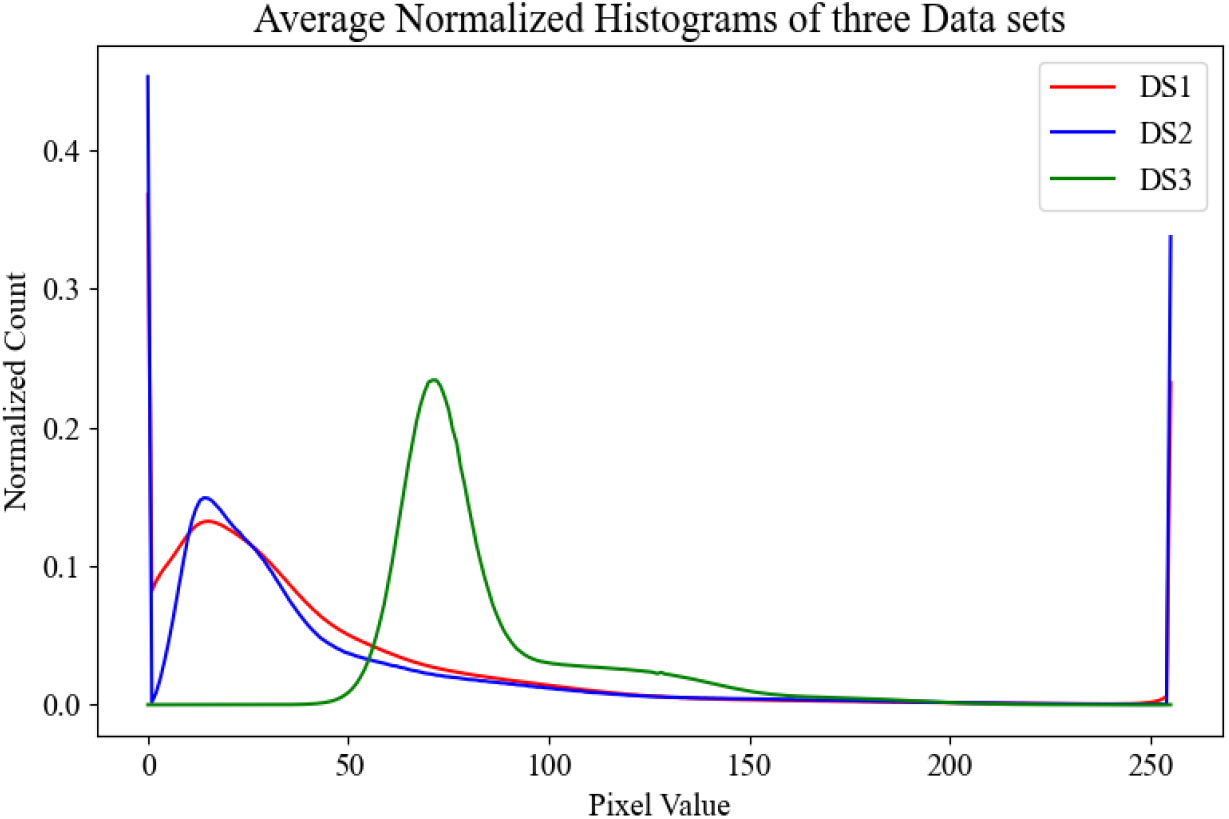
This figure displays the average normalized histograms for each dataset.

### 2.2 Centralized and Local Models

Our hypothesis was that an FL model could be trained to deliver performance on par with a centralized model that has access to all data and would surpass the performance of local models trained solely on locally available data. We established baseline models, both centralized and local, and evaluated their performance against test sets. Local models were trained exclusively on their respective data and then subjected to performance assessments using all three test sets. In contrast, the centralized model underwent training that made use of all three datasets before its evaluation. This rigorous testing methodology provided us with a robust comparative analysis of the performance metrics of these models. Our model’s structure is designed with two main components: an encoder and a classification head (Figure 3). After evaluating various options such as residual network (ResNet), vision transformers (ViT), VGG16, InceptionV3, and EfficientNet, we settled on ResNet18 with 11.2 million and ViT with 4.8 million parameters as the encoding mechanisms for our models, conducting thorough comparisons of their performances across diverse benchmarks. The ViT encoders consist of six transformer blocks and eight heads in the multi-head attention layer. We utilized the Area Under the Receiver Operating Characteristics (ROC) Curve (AUC) as our metric for evaluation. Initial findings indicated that the Adaptive Momentum Estimation with Weight Decay (AdamW) surpassed the Stochastic Gradient Descent (SGD) in terms of performance at both the local and centralized levels, after hyperparameter optimization through grid search. The optimal hyperparameter combination was determined through the maximization of the AUC on the validation set, taking care to prevent data leakage from the test set. To examine the impact of the number of epochs (E) on the models, we trained the models with *E* = 10 and *E* = 100, implementing early stopping based on the AUC of the validation set and patience of ten epochs when *E* = 100. DS1 was processed on a setup equipped with 2*×* Nvidia RTX A6000 graphics cards (node 1), while DS2 was allocated to a machine furnished with 2*×* Nvidia Titan V units (node 2), and DS3 was trained on a node with 8*×* Nvidia GTX1080Ti cards (node 3). Centralized benchmarks were executed on node 3. The summarized results can be found in Table 2.

**Figure 3:**
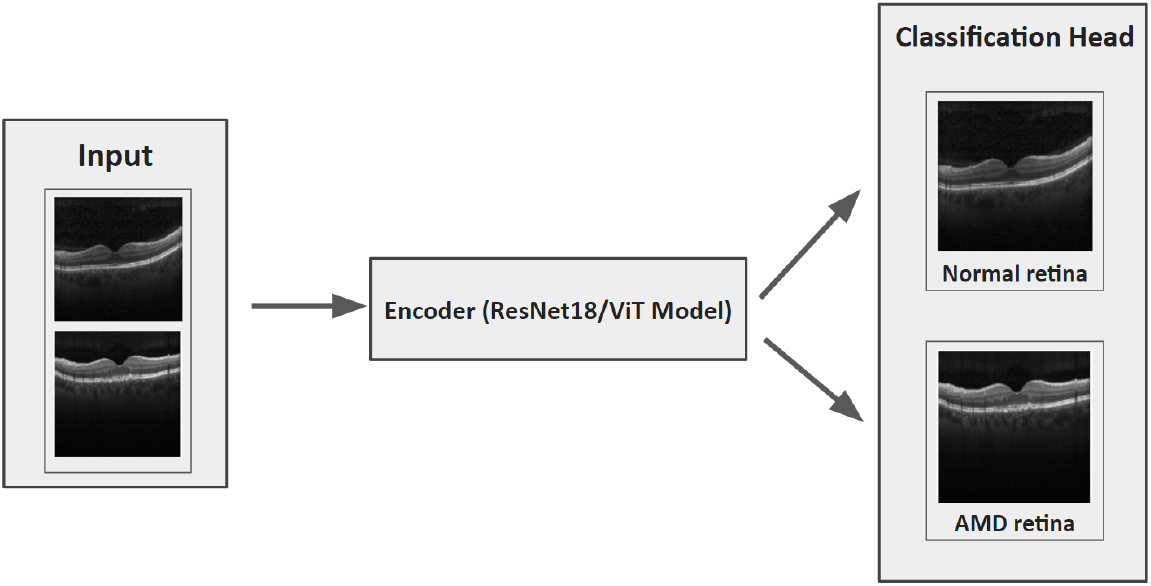
The architecture of our DL model.

**Table 2:** The AUC of the Centralized and Local Models using *E* = 10 and *E* = 100 with early stopping on the validation AUC.

| Method | Test Set |  |  |
| --- | --- | --- | --- |
|  | DS1 | DS2 | DS3 |
| E = 10 |  |  |  |
| Centralized (ResNet18) | 92.78 ± 1.31 | **98.9 ± 0.64** | 97.72 ± 1.5 |
| Centralized (ViT) | 85.64 ± 1.34 | 98.19 ± 0.59 | **99.22 ± 0.38** |
| Local DS1 (ResNet18) | **93.7 ± 0.94** | 98.83 ± 0.63 | 54.14 ± 2.68 |
| Local DS1 (ViT) | 85.23 ± 0.87 | 96.76 ± 0.66 | 88.08 ± 2.27 |
| Local DS2 (ResNet18) | 55.6 ± 1.86 | 80.4 ± 7.21 | 50 ± 0.0 |
| Local DS2 (ViT) | 51.07 ± 0.78 | 75.03 ± 3.96 | 53.63 ± 1.95 |
| Local DS3 (ResNet18) | 49.84 ± 0.56 | 54.65 ± 4.52 | 94.5 ± 2.67 |
| Local DS3 (ViT) | 48.39 ± 1.62 | 56.73 ± 6.53 | 86.3 ± 5.18 |
| E = 100 |  |  |  |
| Centralized (ResNet18) | **94.58 ± 0.62** | 99.05 ± 0.4 | 98.91 ± 0.67 |
| Centralized (ViT) | 87.85 ± 1.2 | **99.18 ± 0.55** | **99.11 ± 0.39** |
| Local DS1 (ResNet18) | 93.97 ± 0.69 | 98.26 ± 0.87 | 51.74 ± 0.66 |
| Local DS1 (ViT) | 88.31 ± 1.5 | 97.7 ± 0.59 | 88.8 ± 2.74 |
| Local DS2 (ResNet18) | 57.03 ± 2.42 | 84.47 ± 8.1 | 50 ± 0 |
| Local DS2 (ViT) | 52.61 ± 1.39 | 83.16 ± 2.84 | 56.6 ± 3.18 |
| Local DS3 (ResNet18) | 49.99 ± 0.01 | 50 ± 0.0 | 91.98 ± 4.02 |
| Local DS3 (ViT) | 48.45 ± 2.83 | 55.49 ± 7.54 | 84.59 ± 4.19 |

### 2.3 FL Framework

Traditional FL algorithms involve a central server that oversees model updates and circulates the global model to all participating nodes. Local models are trained on the respective data and subsequently transmitted back to the server, where they are integrated into the global model [19]. The primary FL algorithms used are FedAvg [21] and Federated Stochastic Gradient Descent [29], as well as their variations.

However, the decentralized character of FL introduces substantial challenges, especially in terms of data heterogeneity and distribution shifts. For instance, in ophthalmology, considerable variations in retinal images across different institutions can be attributable to factors such as the use of distinct imaging devices [6], heterogeneous patient populations [34], and inconsistencies in image acquisition protocols [16].

Addressing these challenges necessitates domain alignment, also referred to as DA. This essential process modifies an ML model trained on one domain to perform proficiently on a related domain. Numerous techniques have been proposed to mitigate the domain shift problem, making it crucial to implement these methods for successful DA. In our FL framework, we have compared four DA strategies alongside FedAvg: FedProx, FedSR, FedMRI, and APFL.

**FedProx**: FedProx [20], is specifically designed to counter the data heterogeneity challenge in FL. It utilizes proximal regularization to incorporate a penalty term into the loss function and avoid overfitting. By maintaining local updates close to the initial global model parameters, FedProx is particularly useful when dealing with not independent and identically distributed data. This ensures each local model does not veer too far from the global model during training, yielding a more resilient global model that performs well across a broader spectrum of data distributions.

**FedSR**: FedSR [23] simplifies the model’s representation and encourages it to extract only essential information. This method employs two regularizers: an L-2 norm regularizer on the representation and conditional mutual information between the data and the representation given by the label. These regularizers limit the quantity of information the representation can contain. By enforcing these regularizers, FedSR facilitates learning data representations that generalize well across diverse domains, all while maintaining data privacy between nodes—a crucial advantage in an FL context.

**FedMRI**: FedMRI [12] addresses the issue of domain shift that might surface during local node optimization. It does so through the implementation of a weighted contrastive regularization, which helps guide the update direction of the network parameters, thus directly rectifying any discrepancies between the local nodes and the server during optimization. This approach contrasts with traditional contrastive learning, which relies on identifying positive and negative pairs from data. In experiments involving multi-institutional data, FedMRI has demonstrated superior performance in image reconstruction tasks compared to state-of-the-art FL methods. As our task resided within the realm of binary image classification, we customized the FedMRI approach. Specifically, we excluded the decoder component and employed the weighted contrastive loss as an auxiliary loss exclusively.

**APFL**: The goal of APFL [10] is to improve the overall performance of a model in an FL setup by considering the distinct data distribution of each participating node. This approach ensures data privacy and model customization. APFL achieves this by adding a level of personalization to the learning process. It involves learning a global model that every node shares, as well as a personalized model that caters to each node’s unique data distribution. The global model identifies common patterns across all nodes, and the personalized model learns from node-specific patterns.

Our FL structure integrated three FL nodes with a central server, and it was developed based on the Flower framework [2]. Before running the local training on these nodes, the server needed to be operational, necessitating the selection of a particular FL strategy, federated, and training configurations. The strategy oversaw several elements of the training and evaluation protocol, such as weight initialization and aggregation. Federated configuration outlined necessary parameters for FL training, encompassing the minimum number of FL nodes needed for training and subsequent evaluation. Further, the training configuration encapsulated requisite parameters for DL model training, including the number of epochs, learning rate, and weight decay.

The procedure to train the FL model generally follows these steps (demonstrated in Figure 4): Initially, the FL strategy (options include FedAvg, FedSR, FedProx, FedMRI, and APFL) will be designated, as well as the FL and training configurations. Subsequently, the server waits for the necessary minimum number of FL nodes to establish a connection, following which it dispatches the training configuration and the initial weights (based on the selected FL strategy) to each node. Each node then updates its local model using the weights received from the server, instigating the training process. Upon completion of the training, each node transmits its local model weights back to the server. Finally, the server aggregates these weights using the designated strategy (such as FedAvg) and reciprocates by sending the updated weights back to each client, marking the conclusion of one round (*R*).

**Figure 4:**
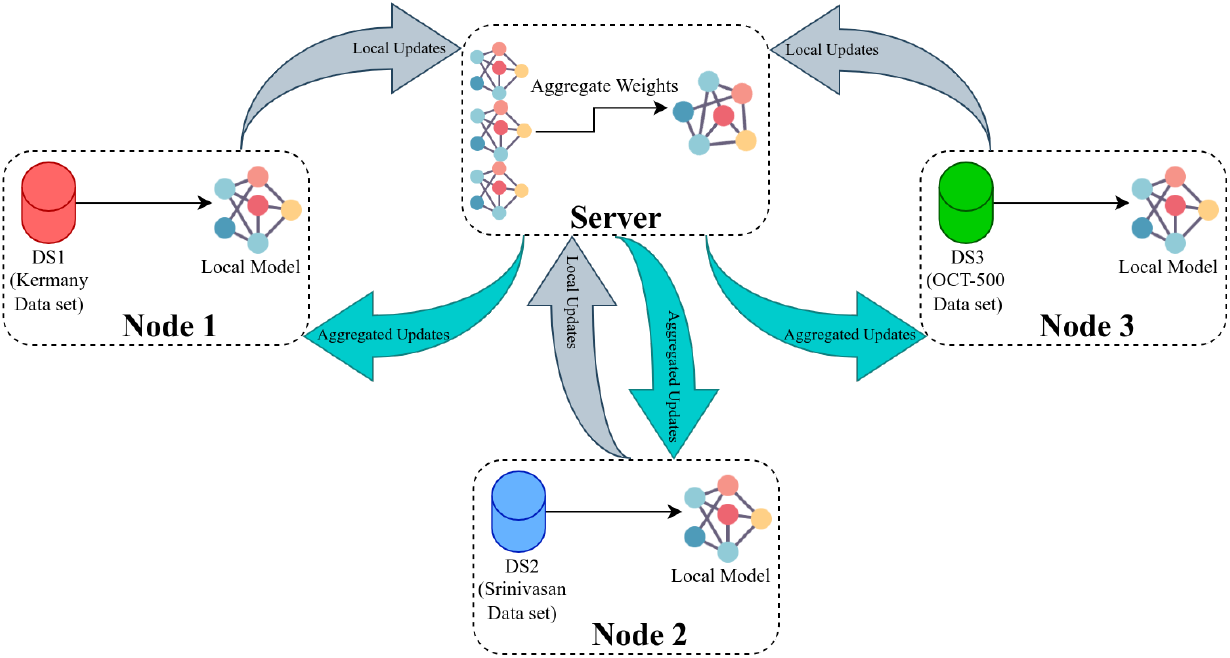
General Overview of FL Framework.

In this framework, there is an optional step called evaluation, where each local node assesses its performance after receiving the global FL model and evaluation configuration over local test sets. The evaluation configuration is similar to the training configuration and may contain various hyperparameters for model evaluation, such as batch size. After evaluation, the performance of each node is sent to the server to demonstrate the FL model’s overall performance across all node test sets. Data heterogeneity can be handled by varying the number of local training epochs. This way, each round of training can be more productive, reducing convergence time and communication costs [22]. To assess the training productivity of each strategy, we examined its AUC with three different allotted local training epochs per round in Table 3.

**Table 3:**
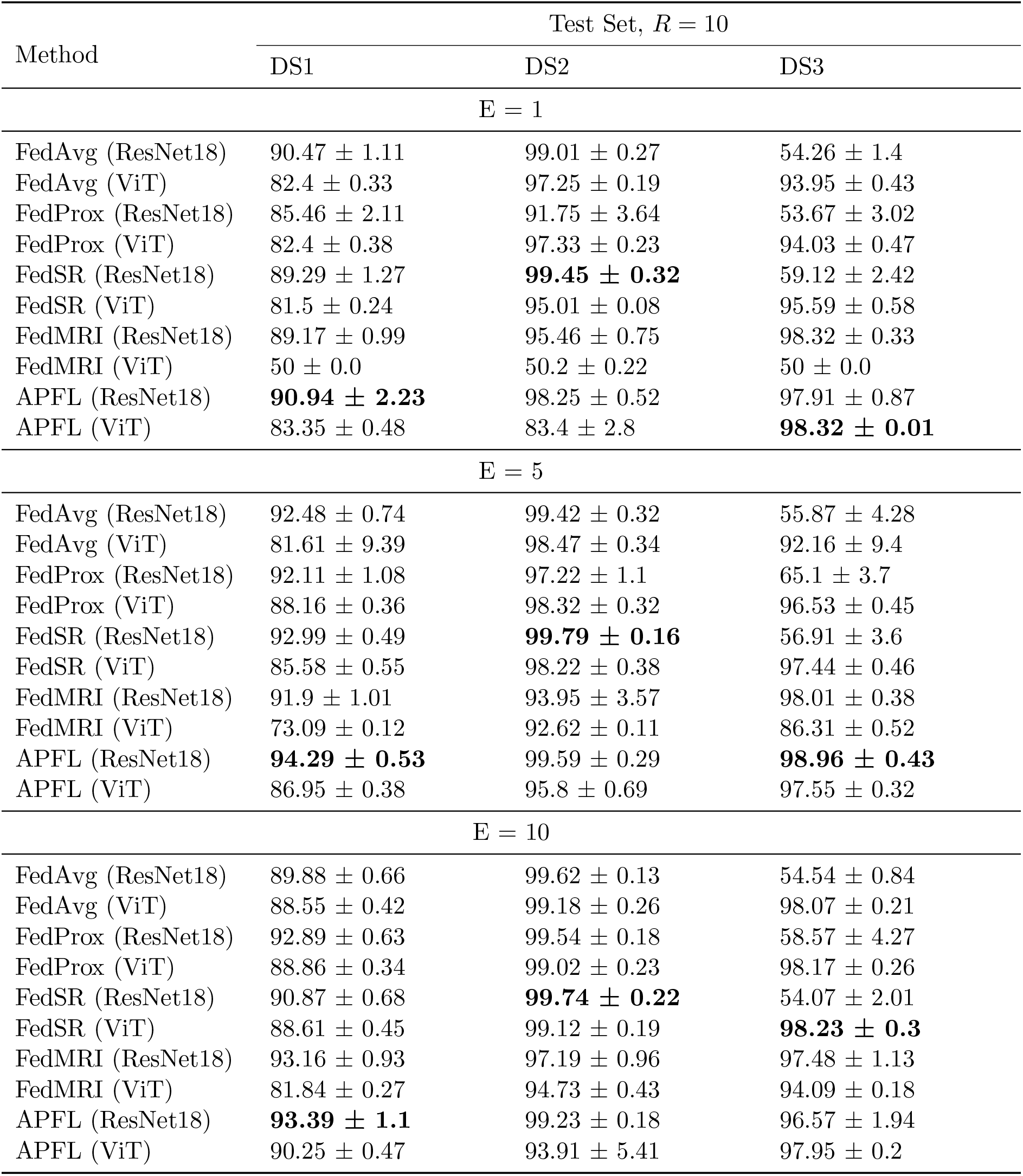
The AUCs of the different FL methods on the three test sets with *E* = 1, *E* = 5, *E* = 10, and *R* = 10.

| Method | Test Set, $R = 10$ | | |
| --- | --- | --- | --- |
|  | DS1 | DS2 | DS3 |
| E = 1 |  |  |  |
| FedAvg (ResNet18) | 90.47 ± 1.11 | 99.01 ± 0.27 | 54.26 ± 1.4 |
| FedAvg (ViT) | 82.4 ± 0.33 | 97.25 ± 0.19 | 93.95 ± 0.43 |
| FedProx (ResNet18) | 85.46 ± 2.11 | 91.75 ± 3.64 | 53.67 ± 3.02 |
| FedProx (ViT) | 82.4 ± 0.38 | 97.33 ± 0.23 | 94.03 ± 0.47 |
| FedSR (ResNet18) | 89.29 ± 1.27 | **99.45 ± 0.32** | 59.12 ± 2.42 |
| FedSR (ViT) | 81.5 ± 0.24 | 95.01 ± 0.08 | 95.59 ± 0.58 |
| FedMRI (ResNet18) | 89.17 ± 0.99 | 95.46 ± 0.75 | 98.32 ± 0.33 |
| FedMRI (ViT) | 50 ± 0.0 | 50.2 ± 0.22 | 50 ± 0.0 |
| APFL (ResNet18) | **90.94 ± 2.23** | 98.25 ± 0.52 | 97.91 ± 0.87 |
| APFL (ViT) | 83.35 ± 0.48 | 83.4 ± 2.8 | **98.32 ± 0.01** |
| E = 5 |  |  |  |
| FedAvg (ResNet18) | 92.48 ± 0.74 | 99.42 ± 0.32 | 55.87 ± 4.28 |
| FedAvg (ViT) | 81.61 ± 9.39 | 98.47 ± 0.34 | 92.16 ± 9.4 |
| FedProx (ResNet18) | 92.11 ± 1.08 | 97.22 ± 1.1 | 65.1 ± 3.7 |
| FedProx (ViT) | 88.16 ± 0.36 | 98.32 ± 0.32 | 96.53 ± 0.45 |
| FedSR (ResNet18) | 92.99 ± 0.49 | **99.79 ± 0.16** | 56.91 ± 3.6 |
| FedSR (ViT) | 85.58 ± 0.55 | 98.22 ± 0.38 | 97.44 ± 0.46 |
| FedMRI (ResNet18) | 91.9 ± 1.01 | 93.95 ± 3.57 | 98.01 ± 0.38 |
| FedMRI (ViT) | 73.09 ± 0.12 | 92.62 ± 0.11 | 86.31 ± 0.52 |
| APFL (ResNet18) | **94.29 ± 0.53** | 99.59 ± 0.29 | **98.96 ± 0.43** |
| APFL (ViT) | 86.95 ± 0.38 | 95.8 ± 0.69 | 97.55 ± 0.32 |
| E = 10 |  |  |  |
| FedAvg (ResNet18) | 89.88 ± 0.66 | 99.62 ± 0.13 | 54.54 ± 0.84 |
| FedAvg (ViT) | 88.55 ± 0.42 | 99.18 ± 0.26 | 98.07 ± 0.21 |
| FedProx (ResNet18) | 92.89 ± 0.63 | 99.54 ± 0.18 | 58.57 ± 4.27 |
| FedProx (ViT) | 88.86 ± 0.34 | 99.02 ± 0.23 | 98.17 ± 0.26 |
| FedSR (ResNet18) | 90.87 ± 0.68 | **99.74 ± 0.22** | 54.07 ± 2.01 |
| FedSR (ViT) | 88.61 ± 0.45 | 99.12 ± 0.19 | **98.23 ± 0.3** |
| FedMRI (ResNet18) | 93.16 ± 0.93 | 97.19 ± 0.96 | 97.48 ± 1.13 |
| FedMRI (ViT) | 81.84 ± 0.27 | 94.73 ± 0.43 | 94.09 ± 0.18 |
| APFL (ResNet18) | **93.39 ± 1.1** | 99.23 ± 0.18 | 96.57 ± 1.94 |
| APFL (ViT) | 90.25 ± 0.47 | 93.91 ± 5.41 | 97.95 ± 0.2 |

During each benchmarking session, one of the nodes played a dual role by serving as both a server and an FL node. The other two resources solely functioned as FL nodes and communicated with the server. Whenever *E* = 10, node 1 assumed the role of both server and FL node. When *E* = 5, node 2 became the server, and when *E* = 1, node 3 took on the role of server. The DL models at each local node were trained using the hyperparameters detailed in the preceding section. Note that, the value of *R* in all FL benchmarks is 10. The hyperparameters, input size, and image transformation have been applied as previously mentioned.

## 3 Results

The summary of outcomes from training a variety of local and centralized models is given in Table 2. These models are evaluated against three distinct test sets at the end of the training phase. The training process employed both ResNet18 and ViT encoders, and the table presents the corresponding performance metrics for each. In the latter part of Table 2, outcomes from training models at *E* = 100 are particularly highlighted. At *E* = 10, the local DS1 ResNet18 achieved superior performance on its native test set, while the centralized ResNet18 and ViT excelled on DS2 and DS3 test sets, respectively. With *E* = 100, centralized models topped the performance charts, with the ResNet18 encoder recording the highest accuracy rates of %94.58 *±* 0.62 on DS1, and the ViT encoder reaching %98.18 *±* 0.55 and %99.11 *±* 0.39 on DS2 and DS3 test sets, respectively.

Moreover, FL strategies such as FedAvg, FedProx, FedSR, FedMRI, and APFL have been meticulously detailed in Table 3. These strategies have been examined in tandem with the employment of ResNet18 and ViT encoders, with the models being trained at *E* = 1, *E* = 5, and *E* = 10. To facilitate easier comprehension, the table specifically highlights in bold the highest AUC for each *E* value. Remarkably, a pattern emerges in the performance of the models on different test sets. The APFL ResNet18 performed the best for DS1. The FedSR ResNet18 showed superior performance for DS2. As for DS3, the APFL ViT, APFL ResNet18, and FedSR ViT performed the best at *E* = 1, *E* = 5, and *E* = 10 respectively. However, it is crucial to bear in mind that the optimal model should maintain a balanced performance across all test sets, and not merely excel in a single one. To ensure consistency, the parameter *R* has been maintained at a constant value of 10 throughout all the testing scenarios.

Figure 5 provides essential information on the performance of the centralized ResNet18 and ViT models across the three test sets at *E* = 100, with the patient parameter set to ten. It also features the exceptional performance of the APFL strategy, denoting it as the leading FL method in this problem.

**Figure 5:**
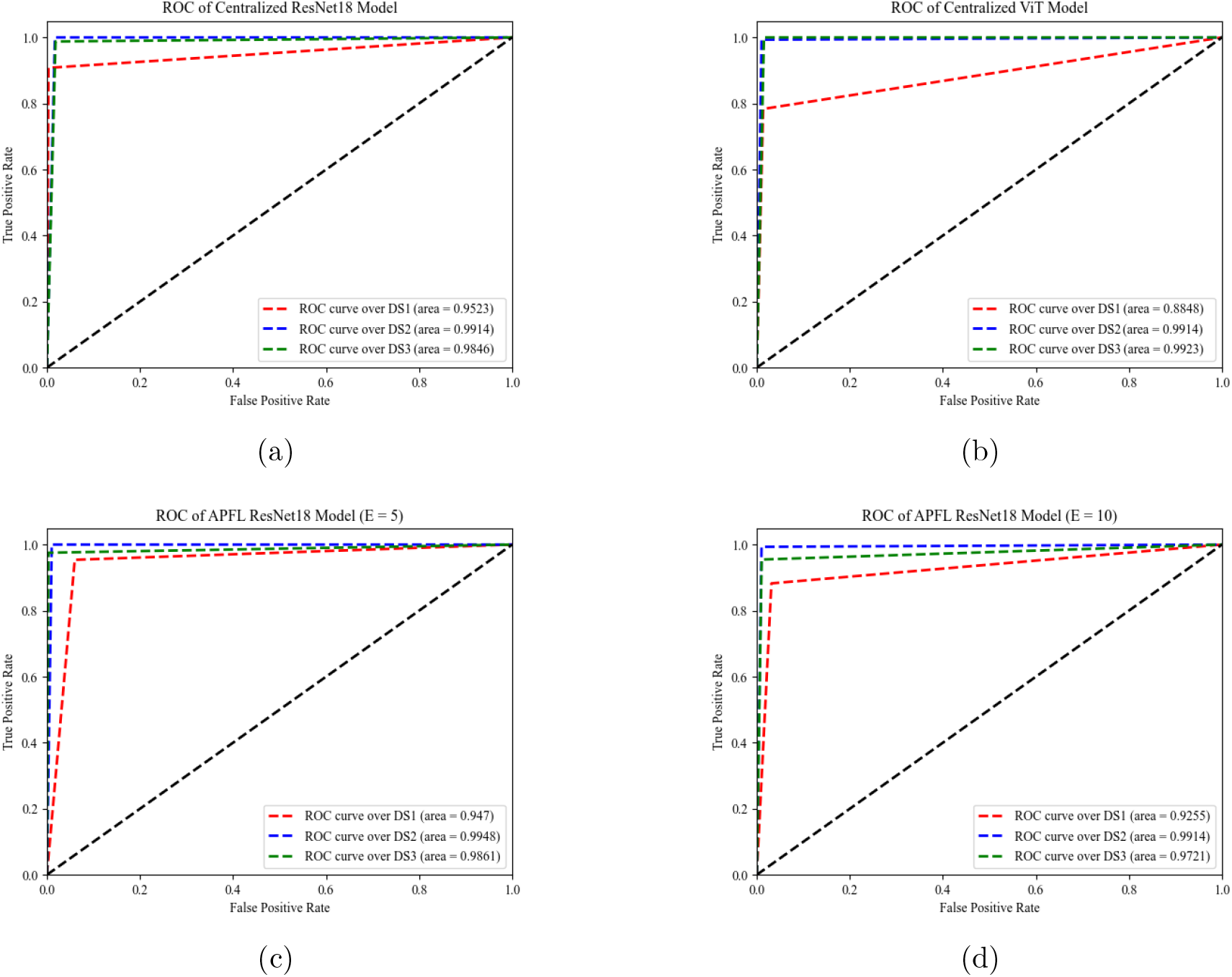
The AUC of centralized models across three datasets, as depicted by ResNet18 and ViT encoders, are illustrated in (**A**) and (**B**), respectively. The same performance measurements, but this time using the APFL strategy with *E* = 5 and *E* = 10, are showcased in (**C**) and (**D**) correspondingly.

Figure 6 depicts the training duration for each model, with a noticeable pattern of longer training times for ViT models in comparison to the ResNet18 equivalents. This trend is consistently apparent across local, centralized, and FL models, even persisting through FL training iterations at *E* = 5 and *E* = 10. The time difference is minimal when training the local model using DS2 - ResNet18 takes about 4-6 seconds, while ViT requires around 5-7 seconds. However, this difference grows when it comes to centralized and FL models, extending up to approximately 40 seconds for training one epoch. Keep in mind that the duration to train an FL model for one epoch is timed from the instant the server dispatches the initial weights to all nodes until it receives and aggregates all the parameters (FL training time). This calculation does not include the time spent on initializing the server, starting the nodes, connecting them to the server, and the evaluation stages. Due to this reason, FL strategies, with the exception of FedSR, tend to take less time to train than centralized models at *E* = 1. Notably, FedSR stands out as having the lengthiest training time among all the benchmarks.

**Figure 6:**
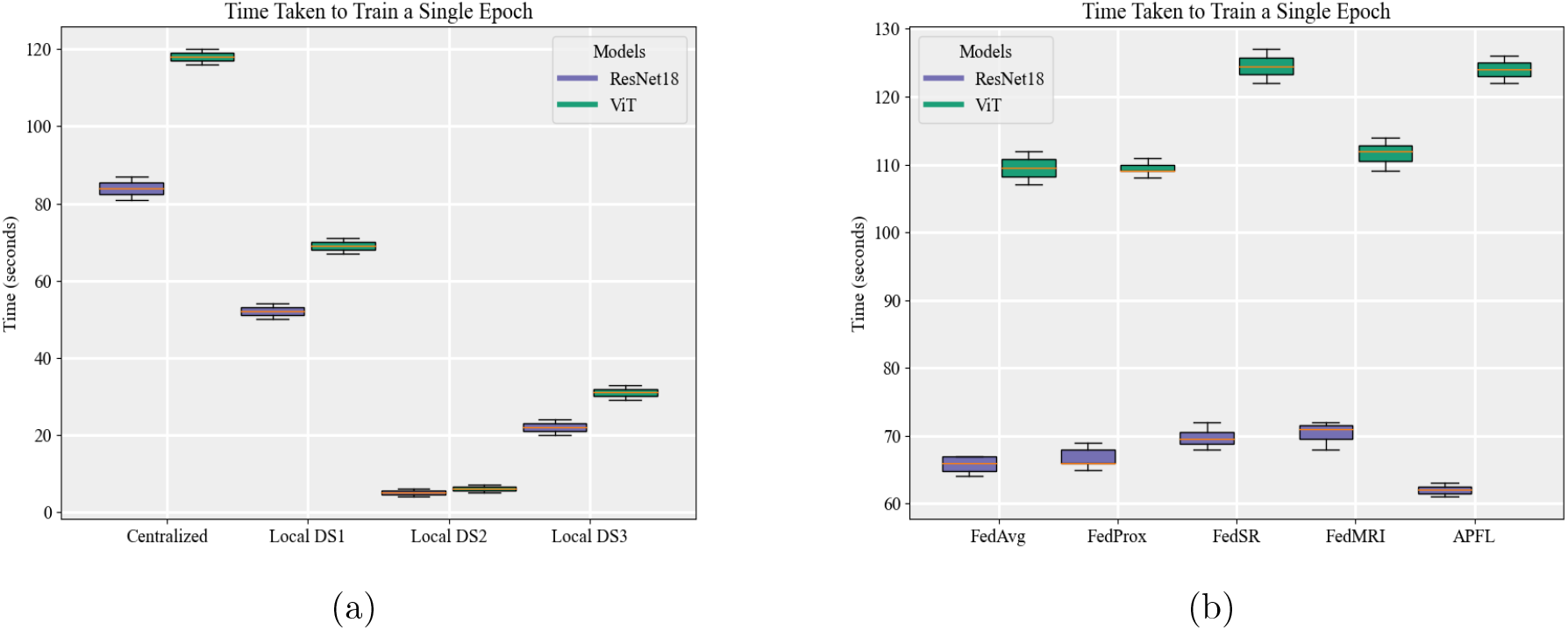
A comparison of the time taken to train an epoch for (**A**) local and centralized models, and (**B**) different FL strategies when local *E* = 1.

This study presented a comprehensive series of experiments exploring the comparative effectiveness of deploying DL models using local, centralized, and FL methodologies across three distinct datasets. The primary focus was the classification of OCT images into Normal and AMD binary categories, for which we utilized ResNet18 and ViT encoders. We also integrated four unique DA methods into our FL strategy to tackle the prevalent issue of domain shift. As our results show, DA FL strategies demonstrated impressive proficiency in training a global model well-suited to this specific problem, achieving competitive performance metrics in comparison to centralized ResNet18 and ViT models despite a lack of access to the entire dataset. These findings underscore the critical role of FL in healthcare settings, where data accessibility is often compromised due to feasibility issues and privacy concerns. By assuring patient confidentiality and facilitating significant insights from distributed learning, FL reinforces its importance in the future of healthcare analytics.

ResNet architectures were selected due to their well-documented efficacy in image classification tasks, the depth of their architectural design enabling complex data pattern learning, and the convenience of existing pre-trained models. ViT were chosen for their ability to incorporate global image context, an invaluable trait for enhancing image classification performance, and their architecture, which eliminates the need for task-specific designs, permitting complex pattern discernment without requiring specialized configurations.

Our experimental process for local and centralized DL models involved two distinct training scenarios: short-duration training over 10 epochs and extended training over 100 epochs. Our intent was to pinpoint the model offering optimal performance when trained over 100 epochs with matching training data, using the validation AUC as the stopping criteria. We also scrutinized the effect of altering the number of local epochs on the training efficiency of the FL strategies, setting *E* values at 1, 5, and 10.

Our study exhibited the anticipated dominance of centralized models over their local counterparts, a result driven by their unrestricted access to all data during the training phase, particularly when *E* is set to its maximum value of 100. A striking observation that emerged during this comparison involved the inconsistent performance of the local DS1 ResNet18 model across different test sets. Although this model exhibited substantial efficacy on its native and DS2 test sets, it encountered difficulties with DS3. This challenge emanated from the disparity in brightness distribution between DS1 and DS2, and DS3, as visualized in Figure 2. A further examination of the performance of the counterpart model, local DS1 ViT, underscored the inherent strength of the ViT architecture’s focus on global features, contributing to its commendable performance (%88.08 *±* 3.17). However, when reviewing the local DS3 and DS2 models, it was evident that they struggled to deliver high-quality results on tests outside their respective training environments. The limited capacity for model generalization (in the case of DS3) and insufficient training data (for DS2) are potential attributing factors. Intriguingly, local ResNet18 models outpaced their ViT counterparts in performance on corresponding test sets. This observation likely stems from the depth and richness in parameters of the ResNet18 architecture, lending it an edge over the ViT models. Besides, the capacity of ViTs to decode complex patterns in the data is amplified with the increase in the volume of available data [11].

Examining the FL training duration, the theoretical expectation would be for parallel training (inherent to FL models) to take less time than sequential training. While we observed a slight reduction in the training time for a single epoch of an FL model, the varying sizes of the datasets (with DS1 being larger) prevented significant outpacing of the centralized models. Training discrepancies between nodes also created bottlenecks, with nodes 2 and 3 having to wait for node 1’s completion. This problem was exacerbated as the *E* value increased, resulting in extended idle periods for quicker nodes. This phenomenon is unique to the training phase; during inference, all nodes use the same model, ensuring consistent inference times.

In the scope of FL, the performance of FedAvg, FedProx, and FedSR models with a ResNet18 encoder was observed to be underwhelming on DS3 test set. This discrepancy in performance is rooted in data heterogeneity, which fosters a drift in the learning process. This drift, predominantly aligned with DS1 and DS2, triggers less desirable outcomes when the aggregated FL model is put to the test on DS3. Interestingly, despite the mediocre performance on DS3, the FedSR ResNet18 model was the standout performer across all *E* values on DS2 test set. Conversely, the three strategies counterparts leveraging the ViT encoder consistently achieved above %81 performance across all test sets. This juxtaposition underscores the potential benefits of employing ViTs over ResNet18, given their intrinsic focus on global features. The use of FedMRI brings a different perspective to the situation. FedMRI ResNet18 showcased promising results across all test sets, even as its counterpart ViT model fared poorly at *E* = 1 and was relatively lackluster at *E* = 5. This highlights the need for more intricate hyperparameter tuning to determine the optimal weighting for contrastive loss. In contrast, the APFL strategy emerged as an exceptional FL approach, consistently yielding an AUC performance exceeding %83 across all tests, irrespective of the encoder used. Remarkably, the APFL ResNet18 model delivered impressive results, nearly matching, and in some instances exceeding, the performance of centralized models. For instance, on DS1 test set, the APFL ResNet18 model achieved an AUC score of %94.29 *±* 0.53 at *E* = 5, closely trailing the %94.58 *±* 0.62 recorded by the centralized ResNet18 model at *E* = 100. On DS2, the model managed to score %99.59 *±* 0.29, surpassing the centralized ViT’s %99.18 *±* 0.55 at *E* = 10. Similarly, on DS3 test set, this model delivered a highly competitive performance of %98.96 *±* 0.43, narrowly trailing the centralized ViT model’s score of %99.22 *±* 0.38 at *E* = 10.

The success of the APFL approach can be attributed to the personalized layer that learns from node-specific data distributions, thereby providing robust and reliable performance. This demonstrates the potential competitiveness of FL models when compared with their centralized counterparts, and indeed, their significant superiority over leading local models. In conclusion, our investigation emphasizes the potential of FL strategies, especially those equipped with adaptive personalization, in developing robust models that deliver consistent performance across varied datasets. This bodes well for future applications in contexts where data privacy and decentralization are of paramount importance.

Our study has several limitations in the training phase, as we opted for a relatively straightforward DL architecture and an aggregation policy based solely on a weighted average. We plan to explore more sophisticated aggregation policies in our future work. Despite these limitations, our findings offer valuable insights into the comparative effectiveness of training simpler architectures for image classification tasks and contribute to our broader understanding of FL strategies. We believe investigating other FL strategies in the future would enrich our understanding of these models’ performance nuances. We also consider the separate classification head as a prospective area of focus, with intelligent weight aggregation policy and normalization of amplitude possibly enhancing FL network performance [13]. Finally, the exploration of deeper models like ResNet50 or ResNet101, or ViTs with more transformer blocks and deeper multi-layer perceptron layers, may alter the performance dynamics and offer fresh insights.

## 4 Discussion

This study presented a comprehensive series of experiments exploring the comparative effectiveness of deploying DL models using local, centralized, and FL methodologies across three distinct datasets. The primary focus was the classification of OCT images into Normal and AMD binary categories, for which we utilized ResNet18 and ViT encoders. We also integrated four unique DA methods into our FL strategy to tackle the prevalent issue of domain shift. As our results show, DA FL strategies demonstrated impressive proficiency in training a global model well-suited to this specific problem, achieving competitive performance metrics in comparison to centralized ResNet18 and ViT models despite a lack of access to the entire dataset. These findings underscore the critical role of FL in healthcare settings, where data accessibility is often compromised due to feasibility issues and privacy concerns. By assuring patient confidentiality and facilitating significant insights from distributed learning, FL reinforces its importance in the future of healthcare analytics.

ResNet architectures were selected due to their well-documented efficacy in image classification tasks, the depth of their architectural design enabling complex data pattern learning, and the convenience of existing pre-trained models. ViT were chosen for their ability to incorporate global image context, an invaluable trait for enhancing image classification performance, and their architecture, which eliminates the need for task-specific designs, permitting complex pattern discernment without requiring specialized configurations.

Our experimental process for local and centralized DL models involved two distinct training scenarios: short-duration training over 10 epochs and extended training over 100 epochs. Our intent was to pinpoint the model offering optimal performance when trained over 100 epochs with matching training data, using the validation AUC as the stopping criteria. We also scrutinized the effect of altering the number of local epochs on the training efficiency of the FL strategies, setting *E* values at 1, 5, and 10.

Our study exhibited the anticipated dominance of centralized models over their local counterparts, a result driven by their unrestricted access to all data during the training phase, particularly when *E* is set to its maximum value of 100. A striking observation that emerged during this comparison involved the inconsistent performance of the local DS1 ResNet18 model across different test sets. Although this model exhibited substantial efficacy on its native and DS2 test sets, it encountered difficulties with DS3. This challenge emanated from the disparity in brightness distribution between DS1 and DS2, and DS3, as visualized in Figure 2. A further examination of the performance of the counterpart model, local DS1 ViT, underscored the inherent strength of the ViT architecture’s focus on global features, contributing to its commendable performance (%88.08 *±* 3.17). However, when reviewing the local DS3 and DS2 models, it was evident that they struggled to deliver high-quality results on tests outside their respective training environments. The limited capacity for model generalization (in the case of DS3) and insufficient training data (for DS2) are potential attributing factors. Intriguingly, local ResNet18 models outpaced their ViT counterparts in performance on corresponding test sets. This observation likely stems from the depth and richness in parameters of the ResNet18 architecture, lending it an edge over the ViT models. Besides, the capacity of ViTs to decode complex patterns in the data is amplified with the increase in the volume of available data [11].

Examining the FL training duration, the theoretical expectation would be for parallel training (inherent to FL models) to take less time than sequential training. While we observed a slight reduction in the training time for a single epoch of an FL model, the varying sizes of the datasets (with DS1 being larger) prevented significant outpacing of the centralized models. Training discrepancies between nodes also created bottlenecks, with nodes 2 and 3 having to wait for node 1’s completion. This problem was exacerbated as the *E* value increased, resulting in extended idle periods for quicker nodes. This phenomenon is unique to the training phase; during inference, all nodes use the same model, ensuring consistent inference times.

In the scope of FL, the performance of FedAvg, FedProx, and FedSR models with a ResNet18 encoder was observed to be underwhelming on DS3’s test set. This discrepancy in performance is rooted in data heterogeneity, which fosters a drift in the learning process. This drift, predominantly aligned with DS1 and DS2, triggers less desirable outcomes when the aggregated FL model is put to the test on DS3. Interestingly, despite the mediocre performance on DS3, the FedSR ResNet18 model was the standout performer across all *E* values on DS2’s test set. Conversely, the three strategies counterparts leveraging the ViT encoder consistently achieved above %81 performance across all test sets. This juxtaposition underscores the potential benefits of employing ViTs over ResNet18, given their intrinsic focus on global features. The use of FedMRI brings a different perspective to the situation. FedMRI ResNet18 showcased promising results across all test sets, even as its counterpart ViT model fared poorly at *E* = 1 and was relatively lackluster at *E* = 5. This highlights the need for more intricate hyperparameter tuning to determine the optimal weighting for contrastive loss. In contrast, the APFL strategy emerged as an exceptional FL approach, consistently yielding an AUC performance exceeding %83 across all tests, irrespective of the encoder used. Remarkably, the APFL ResNet18 model delivered impressive results, nearly matching, and in some instances exceeding, the performance of centralized models. For instance, on DS1’s test set, the APFL ResNet18 model achieved an AUC score of %94.29 *±* 0.53 at *E* = 5, closely trailing the %94.58 *±* 0.62 recorded by the centralized ResNet18 model at *E* = 100. On DS2, the model managed to score %99.59 *±* 0.29, surpassing the centralized ViT’s %99.18 *±* 0.55 at *E* = 10. Similarly, on DS3’s test set, this model delivered a highly competitive performance of %98.96 *±* 0.43, narrowly trailing the centralized ViT model’s score of %99.22 *±* 0.38 at *E* = 10.

The success of the APFL approach can be attributed to the personalized layer that learns from node-specific data distributions, thereby providing robust and reliable performance. This demonstrates the potential competitiveness of FL models when compared with their centralized counterparts, and indeed, their significant superiority over leading local models. In conclusion, our investigation emphasizes the potential of FL strategies, especially those equipped with adaptive personalization, in developing robust models that deliver consistent performance across varied datasets. This bodes well for future applications in contexts where data privacy and decentralization are of paramount importance.

Our study has several limitations in the training phase, as we opted for a relatively straightforward DL architecture and an aggregation policy based solely on a weighted average. We plan to explore more sophisticated aggregation policies in our future work. Despite these limitations, our findings offer valuable insights into the comparative effectiveness of training simpler architectures for image classification tasks and contribute to our broader understanding of FL strategies. We believe investigating other FL strategies in the future would enrich our understanding of these models’ performance nuances. We also consider the separate classification head as a prospective area of focus, with intelligent weight aggregation policy and normalization of amplitude possibly enhancing FL network performance [13]. Finally, the exploration of deeper models like ResNet50 or ResNet101, or ViTs with more transformer blocks and deeper multi-layer perceptron architectures, may alter the performance dynamics and offer fresh insights.

## Acknowledgments

We acknowledge funding support from the University of North Carolina at Charlotte Faculty Research Grant (FRG).

## Data Availability Statement

To access the datasets analyzed in this study, use the following links: DS1, DS2, and DS3.

